# Complementary metrics of human auditory nerve function derived from compound action potentials

**DOI:** 10.1101/213157

**Authors:** Kelly C. Harris, Kenneth I. Vaden, Carolyn M. McClaskey, James W. Dias, Judy R. Dubno

## Abstract

Declines in auditory nerve (AN) function contribute to suprathreshold auditory processing and communication deficits in individuals with normal hearing, hearing loss, hyperacusis, and tinnitus. Procedures to characterize AN loss or dysfunction in humans are limited. We report several novel complementary metrics to characterize AN function noninvasively in humans using the compound action potential (CAP), a direct measure of summated AN activity. We examined how these metrics change with stimulus intensity, and interpreted these changes within a framework of known physiological properties of the basilar membrane and AN. Our results reveal how neural synchrony and the recruitment of AN fibers with later first-spike latencies likely contribute to the CAP, affect auditory processing, and differ with noise exposure history in younger adults despite normal pure-tone thresholds. Moving forward, these new metrics, when applied to patient populations, can provide a means to characterize cochlear synaptopathy and other deficits in AN function in humans.

New and noteworthy

Loss or inactivity of auditory nerve (AN) fibers is thought to contribute to suprathreshold auditory processing deficits, but evidence-based methods to assess these effects are not available. We describe several novel metrics that may be used to quantify neural synchrony and characterize AN function.

## Introduction

The healthy human cochlea contains ~30,000 auditory nerve (AN) fibers and represents the sole route from the inner ear to the central auditory system. All acoustic information must be encoded in the spike timing and rates of AN fibers. Studies in laboratory animals have shown that moderate noise exposure and aging can lead to a loss or inactivity of AN fibers (Furman et al. 2013; Kujawa and Liberman 2009; 2015; Schmiedt et al. 1996), especially those with low spontaneous rates (low-SR). Loss or inactivity of low-SR fibers can lead to compensatory changes throughout the auditory system and may disrupt suprathreshold auditory processes (Chambers et al. 2016). Despite a potential impact on suprathreshold processing, losses of up to 80% of AN synapses may not affect pure-tone detection thresholds (Lobarinas et al. 2013; Schuknecht and Woellner 1953). This suggests that standard clinical assessments of human hearing based on detection thresholds (‘audiogram’) may appear normal despite significant AN loss. The loss of AN synapses, termed “cochlear synaptopathy,” has been hypothesized to underlie declines in suprathreshold processing in individuals with normal pure-tone thresholds and other auditory perceptual anomalies including hyperacusis and tinnitus (Hickox and Liberman 2014; Plack et al. 2014; Schaette and McAlpine 2011).

The compound action potential (CAP) is an extracellular potential reflecting the summed response of a population of AN fibers. Here, we present several new metrics developed from the CAP response (Figures 1-2), which may be used individually or in combination to non-invasively characterize AN function and fiber loss in humans. We introduce and validate five metrics derived from the averaged CAP response that exploit known differences in response patterns of high-SR and low-SR fibers. Compared to high-SR fibers, low-SR fibers have higher thresholds, larger dynamic ranges, better preservation of timing information, later first-spike latencies and slower conduction velocities (Bourien et al. 2014; Heil and Irvine 1997; Liberman 1978). Due to the effects of signal averaging, differences in spike timing of individual fibers result in an averaged response that is wider than either the single trial responses or the individual action potentials. The width of this response is dependent on the conduction velocity and fiber diameter of the contributing neurons, which vary depending on the fiber’s SR and location in the cochlea. Fibers with high-SRs and higher characteristic frequencies are known to have earlier first-spike latencies than fibers with low-SRs and lower characteristic frequencies (Heil and Irvine 1997). The onset of an averaged response reflects the earliest single-trial response, or fibers with earlier first-spike latencies, whereas the peak latency and width of the response reflect the contribution of fibers with later first-spike latencies. These relationships form the basis of the classic finding that increasing stimulus intensity results in higher CAP amplitudes and shorter latencies(Thornton 1976). As intensity increases, peripheral auditory filters broaden, which has the effect of recruiting higher frequency fibers located in the cochlear base. With increasing intensity, neural synchrony also increases and the population response of AN fibers has an earlier latency (Figure 3A), resulting in higher CAP amplitudes and shorter latencies. We hypothesize that individual differences in the rate of change in amplitude and peak latency with increasing level may reflect differences in the ability to recruit low-SR fibers. That is, individuals who recruit relatively more fibers with increasing level will have steeper amplitude-intensity functions with broader responses that yield prolonged peak latencies (Figure 3B). We propose that using this multi-metric approach will improve our ability to identify and characterize AN dysfunction in individual listeners.

**Figure 1.**
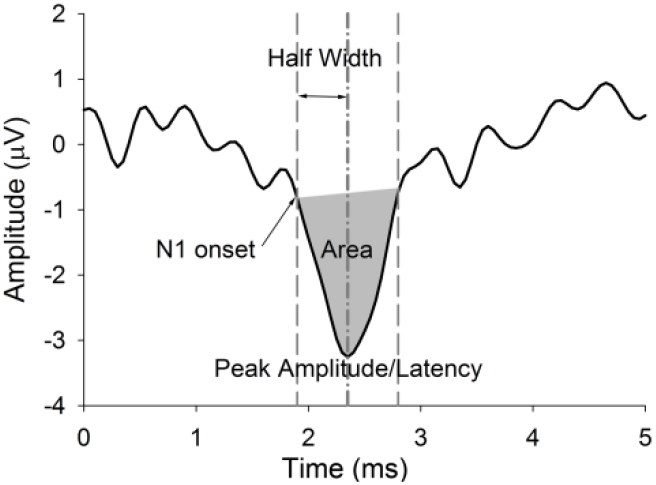
A representative CAP response from one participant and illustration of 5 metrics: ***Peak amplitude***: Peak-to-baseline amplitude calculated in reference to the average baseline (-1 to 1 ms) (µV) ***Peak latency***: Latency of the peak amplitude (ms) ***Onset latency***: 90% fractional peak latency at N1 onset (ms) ***Half-width***: Time from onset latency to peak latency (ms) ***Area***: Numerical integration from N1 onset to offset (gray area in Fig. 1) (µV/ms)

**Figure 2.**
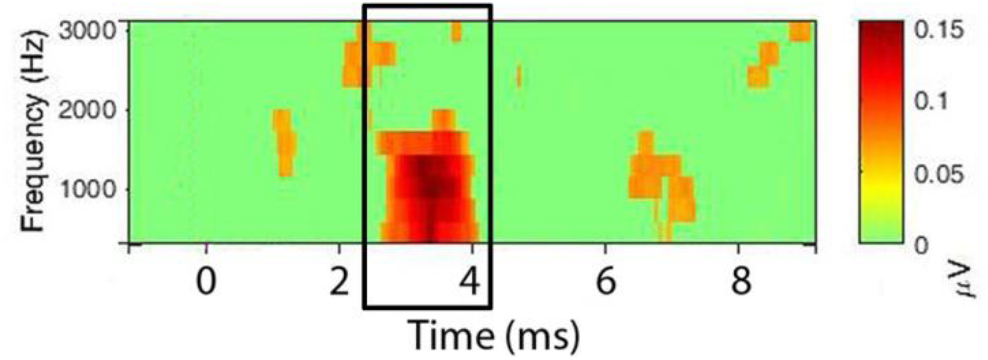
PLV time-frequency plot. Time-frequency analysis was used to estimate PLV of N1 (EEGlab) within linearly spaced frequencies from 625 to 3120 Hz. The x-axes represent time (signal onset = 1 ms). The y-axis represents frequency. Window size was 1.6 ms, with a padratio of 2. PLVs were extracted from a 2 ms window surrounding the peak of N1.

**Figure 3.**
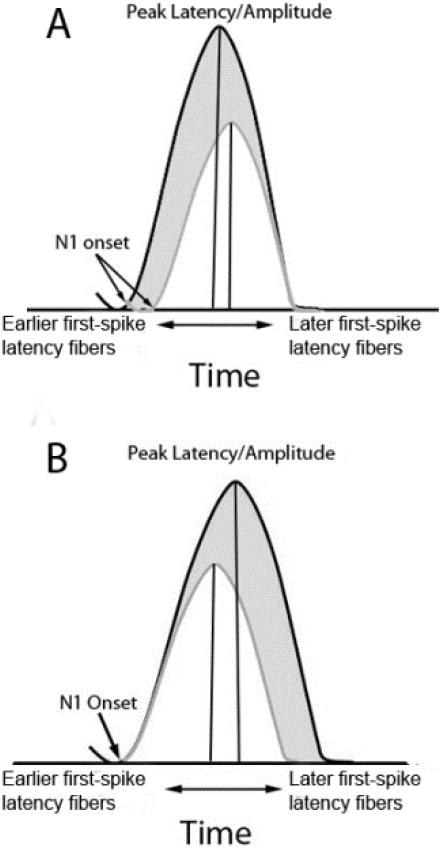
Schematics depicting the association between amplitude and latency in the CAP response. **A.** Schematic depicting the classic changes in the CAP response with increasing level *within a participant* (white to gray), showing shorter onset and peak latencies. As stimulus intensity increases, the probability of individual nerve fibers firing increases; there is also increased neural synchrony and a basal ward spread of excitation along the basilar membrane, resulting in the recruitment of higher frequency AN fibers that have earlier first-spike latencies. Thus, the shift in both the onset of N1 response and the peak of the N1 response reflects a change across the neural population. The increase in amplitude with decreasing latency likely reflects improved synchrony and larger numbers of neurons activated with earlier first-spike latencies (higher frequency). **B**. Schematic depicting hypothesized contribution of slower low-SR fibers to the amplitude and latency of the CAP *across participants in response to a high-level click* (white and gray). The fibers with the earliest first-spike latency contribute to the onset of the response whereas fibers with longer first-spike latencies contribute to the width and peak latency. Accordingly, larger N1 peak amplitudes, prolonged peak latencies, but similar onset times *across participants* are consistent with greater recruitment of AN fibers with more varied and slower first-spike latencies in those participants with the steepest growth of N1 amplitudes (gray response).

In addition to the above mentioned metrics drawn from the time-averaged response we analyzed responses to individual stimuli in the time- frequency domain to quantify changes in neural synchrony across stimulus presentations. Similar to techniques previously used at the level of the brainstem and cortex (Clinard and Tremblay 2013; Harris 2014; Zhu et al. 2013), we estimated the phase locking value (PLV) of the AN. PLV reflects the uniformity of the phase at a specific time and frequency across trials. Neural synchrony across trials is crucial in the generation of an averaged potential. Deficits in neural synchrony are hypothesized to contribute to several functional abnormalities, including age-related declines in auditory processing and auditory neuropathy.

Finally, we integrate these new measures to demonstrate how the recruitment of AN fibers with later first-spike latencies and neural synchrony contribute to the CAP, affect auditory processing, and differ with noise exposure history. The development of objective, non-invasive measures has wide-spread clinical ramifications. A loss or inactivity of AN fibers may disrupt temporal processing and result in undersampling of the sound waveform, which are particularly detrimental to understanding speech in competing noise (Lopez-Poveda 2014).

## Methods

### 2.1 Participants

26 participants, aged 22-30 years, were recruited from the Charleston community [mean age=25.3 years; 15 females]. Participants had pure-tone thresholds ≤20dB HL at 250, 500, 1000, 2000, 3000, 4000 Hz, and 8000 Hz; differences in thresholds between right and left ears did not exceed 15 dB HL at any frequency. Participants also had normal otoscopic findings and tympanometric measures. Participants provided written informed consent before participating in this MUSC Institutional Review Board approved study.

### 2.2 CAP acquisition

Measures were obtained with the participants seated in an acoustically and electrically shielded booth. The CAP was elicited by 100 µS click, rectangular pulses, alternating polarity, presented at 11.1/s in 10-dB steps ranging from 70-110 dB pSPL to the right ear through an insert earphone (ER-3c; Etymotic Technologies). These earphones introduce a 1-ms delay, therefore stimulus presentation occurs at 1 ms, rather than at 0 ms. CAPs were recorded in blocks of 1100 trials (550 of each polarity). Each intensity level was presented twice. Signal level was varied randomly across blocks from 70 to 110 dB SPL, with the exception that the highest level could not occur first.

Electrical signals were recorded with a tympanic membrane electrode (Sanibel, Eden Prairie, MN) as the active electrode, a surface electrode on the contralateral mastoid (inverting), and a low forehead electrode (ground). This montage optimizes activity from the AN, or N1. Participants slept and/or relaxed with their eyes closed during EEG recordings.

EEGs were recorded continuously using a custom head stage (Tucker Davis Technologies, 30x Gain) connected to the bipolar channels of a Neuroscan SynAmpsRT (AC mode, 2010x Gain) and digitized at a rate of 20,000 Hz. EEG data were analyzed offline in Matlab using EEGlab and the ERPlab toolbox. EEG signals were pass-band filtered between 200 and 2500 Hz. The filtered data were then epoched from -1 to 10 ms and baseline corrected to a -1ms to 1 pre-stimulus baseline. Trials were identified and rejected based on a moving window with peak-to-peak threshold detection of 50 µV. Epoched responses for the remaining trials were averaged. An N1 was considered present if it occurred within 2 to 4.5 ms post-stimulus onset, was repeatable (identified in both runs at the same intensity), and identified by two independent and experienced reviewers.

### 2.3 Onset Latency, Peak Latency and Amplitude

In addition to the conventional measure of peak latency of the averaged CAP response, we measured the latency at the *onset* of the CAP response. The CAP peak-to-baseline amplitude and peak latency of the N1 were identified using visual overlay cursors on a computer monitor. Peak amplitude is measured in reference to the average baseline (-1 to 1 ms). Peak selection was performed by two independent reviewers and assessed for repeatability across multiple runs. The onset of the N1 of the CAP was calculated in ERPlab using the fractional peak latency function. The fractional peak latency is calculated from the peak, back in time, as the point at which the signal reaches 10% of the fraction of the peak. Rather than 0%, the 10% point was selected because of the influence of trial-by-trial latency variability, such that the onset time of an averaged evoked response will be substantially earlier than the average onset of the single trials, and the 10% peak amplitude point may be more representative of the average single-trial onset times (Luck 2004). Visually, this time point corresponded well with the “starting point” of the evoked response.

### 2.4 Half width and Area

The half width of the CAP response was calculated as the time (in ms) from the onset of the CAP to the peak latency. We used numerical integration to calculate the area of the CAP with units of µV/ms.

### 2.5 PLV

PLV, also called inter-trial coherence (ITC), is the relative number of responses at a particular phase or time-post onset relative to the stimulus(Delorme et al. 2007). An important advantage of PLV is that it is independent of differences in baseline levels of noise and signal power. In addition, given that it is a measure of neural synchrony, PLV may provide a human parallel to the phase-locking measures acquired in single-unit auditory neurophysiological measures in laboratory animals (Zhu et al. 2013) or simulated with predictions from physiologically inspired auditory-nerve computational models (Zilany et al. 2009). PLVs were measured using time-frequency analysis performed using a continuous Morlet wavelet transform as implemented in EEGlab (Delorme and Makeig 2004). This process is fully described in Delorme and Makeig (2004). Briefly, PLV measures take a value of 0, meaning absence of synchronization between the EEG and the stimulus event, and 1, meaning perfect synchronization. Typically for *n* trials, if F_k_ (f,t) is the spectral estimate of trial *k* at frequency *f* and time *t*.

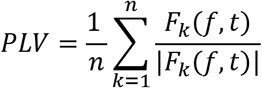

PLV was computed with linearly spaced frequencies from 625 Hz to 3120 Hz, with a padratio of 2, and a window size of 32 samples. PLVs were assessed relative to pre-stimulus baseline (-1 to 1 ms) (*p*<0.001, bootstrap permutation). A single estimate of PLV was obtained for each subject as the median PLV across a 2 ms window surrounding the CAP peak response from 625 to 2500 Hz. A color-coded representation of the PLV data is provided in Figure 3, with PLV images across frequency, where significance levels are assessed using surrogate data by randomly shuffling the single-trial spectral estimates from different baseline latency windows.

### 2.5 Acoustic Reflex Measurements

It has been suggested that CAP measures may be confounded by individual differences in thresholds of the acoustic reflex (Liberman et al. 2016; Valero et al. 2016). To test this hypothesis, acoustic reflex thresholds were measured in a subset of participants (N = 6) using the GSI TympStar (Viasys Health Care, Madison, Wisconsin). By using the external stimuli option, the same hardware used to measure the CAP was used to present a 1.5-s click train at levels ranging from 80 dB pSPL to 120 dB pSPL in 5-dB steps.

### 2.6. Interaural time difference (ITD) Digit Segregation Task

We examined associations between CAP metrics, PLV, and a behavioral temporal processing task. The digit segregation task was previously found to relate to brainstem encoding and hypothesized to reflect differences in AN function (Bharadwaj et al. 2014). Recorded spoken digits of a female speaker were monotonized (184 Hz, close to the natural pitch of the voice). Each trial consisted of two simultaneous sequences of three spoken digits each (containing numbers between 1 and 4). Sequences were differentiated by their ITD. The ITD was used to spatialize the digits to the right or left of midline. A visual cue (light on vote box) and an auditory cue (noise occurring at same ITD as the tokens) was presented 2 s before the onset of the digits, identifying the direction of the target stream (left or right based on ITD). Target direction was randomized on each trial. The ITD size in each trial was drawn uniformly from a set of ITDs of 100, 200, 300, 400, 800, and 1200 µs. Each ITD was presented 210 times. Following the digit sequence presentation, participants indicated the three digits of the target sequence using button presses. Feedback was provided as follows: a green light indicated that all three digits were identified correctly from the target stream; a yellow light indicated that two digits were identified correctly, and a red light indicated that fewer than two digits were identified correctly. The target digit stream was compared to the participant’s response and each digit was recorded as correct or incorrect. The ITD Digit Segregation task was completed by 24 out of 26 participants.

### 2.7 Noise Exposure History

To assess the relationship between self-report noise exposure history and CAP metrics and PLV, participants completed a noise history questionnaire. This questionnaire asks participants to report if they have a positive history for one or more noise exposure categories: (1) noisy work environments, (2) guns, (3) loud music, (4) power tools, (5) farm machinery, and (6) sudden loud noises. Noise history questionnaires were completed by 25 of 26 participants. Based on their responses, participants were divided into two groups; those who answered ‘yes’ to one or more of the noise history questions were included in the “exposed” group(N=15, 8 females) and the remaining participants were included in the “non-exposed” group (N=10, 7 females). Additional details on the noise history questionnaire are included in (Dubno et al. 2013).

### 2.8 Data Analyses

Test-retest reliability of our CAP metrics was assessed using Intraclass Correlation Coefficients (ICCs). Linear regression and repeated measures analysis of variance (ANOVA) were used to assess the extent to which each metric changed with increasing signal intensity. We hypothesized that, due to their higher thresholds and larger dynamic range, the contribution of low-SR AN fibers would be greater at higher but not at lower intensity levels. To test this hypothesis, we examined associations (Pearson product correlations) between metrics at a lower level (80 dB pSPL) and a higher level (110 dB pSPL). Bootstrapping procedures with 10,000 estimates were used to produce 95% confidence intervals for Pearson product correlations. We used linear regression and model testing to assess the independent contribution of our estimates of neural synchrony (PLV) and the contribution of slower AN fibers (peak latency at 110 dB pSPL) to the growth of N1 amplitude. Associations between the N1 response, participants’ hearing thresholds and temporal processing (ITD Digit Segregation Task performance) were modeled using linear mixed-effects regression models, as further explained in the results. Differences in outcomes between the exposed and non-exposed noise history groups were assessed using independent sample t-tests. In order to better visualize the variance across participants and the associations between variables, data are primarily displayed as scatter plots.

## 3. Results

### 3.1 Test-retest reliability

We first analyzed test-retest reliability of the six new metrics (PLV plus five from CAP). Click-evoked CAP recordings showed an N1 present in all 26 participants at levels of 80 dB pSPL and higher. A response was present at 70 dB pSPL in 24 of 26 participants. ICCs ranged from 0.68 to 0.98 for all six metrics across the two runs at the same intensity level (statistically significant ICCs are ≥0.67). Test-retest reliability was also measured across sessions in two participants by re-recording the CAP 1 week after the initial measures. The ICCs from the N1 metrics across sessions were >0.9 at signal levels from 70-110 dB pSPL, suggesting high test-retest reliability across sessions for each CAP metric. The same trained audiologist placed the TM electrode for all participants, likely increasing the repeatability across sessions.

### 3.2 CAP metrics-Intensity functions

We used repeated measures ANOVA to examine the extent to which each metric changed with increasing stimulus intensity. Individual subject data and mean regression lines are provided in Figure 4A-F. With increasing stimulus intensity, peak-to-baseline amplitude significantly increased (became more negative)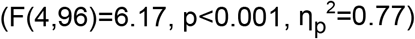, peak latency 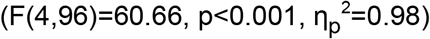 and onset latency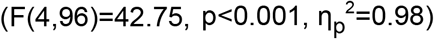 significantly decreased, and the width of the response (half-width) increased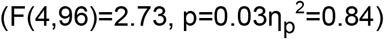. This latency decrease and amplitude increase is consistent with previous reports (Dau 2003; Mehraei et al. 2016). PLV increased with increasing level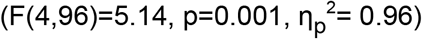, consistent with reports in laboratory animals of improved neural synchrony with increasing intensity(Heil and Irvine 1997). Increased amplitudes with higher level stimuli can be explained by the increased PLV and the recruitment of higher threshold AN fibers. As described earlier, decreased onset and peak latencies are attributed to the broadening of peripheral auditory filters with increasing level. This broadening yields shorter impulse times/earlier first-spike latencies and an excitation pattern that peaks at higher frequencies along the basilar membrane (Harte et al. 2009).Our measure of CAP area was more variable. Although not significant, area showed the general trend of increasing with increasing stimulus intensity (p>0.05).Figure 5 summarizes the multivariate pattern of intensity-dependent changes occurring among N1 peak amplitude, N1 peak latency, N1 half width, and PLV.

**Figure 4.**
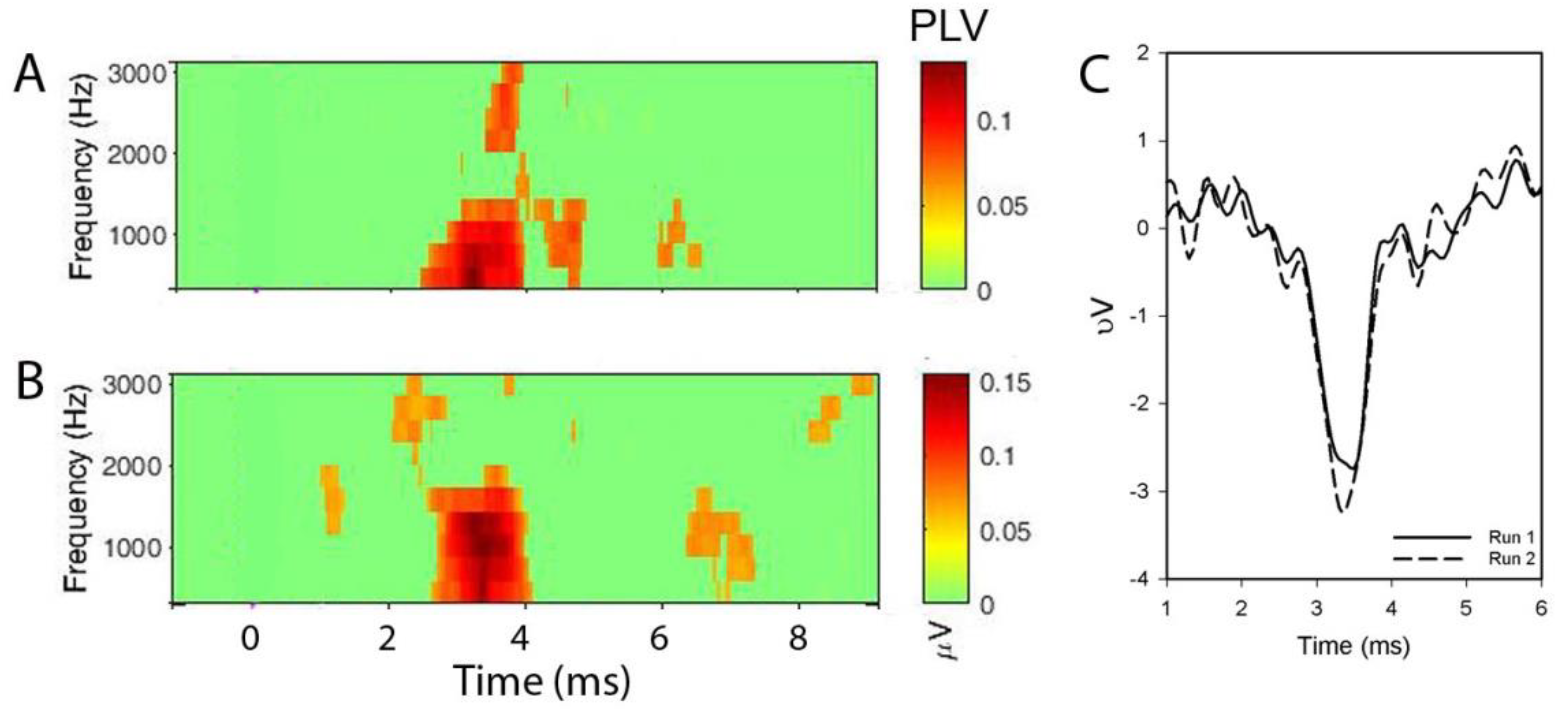
CAP test-retest reliability. CAPs were re-recorded after 1 week from two participants. Two PLV time-frequency plots **(A, B)** and two CAP responses **(C)** recorded one week apart are provided for one of the two participants. The x-axes in Panels A-C represent time (signal onset = 1 ms). The y-axis in Panels A-B represents frequency. Test-retest reliability was estimated using Intraclass Correlation Coefficients (ICC >0.9).

**Figure 5.**
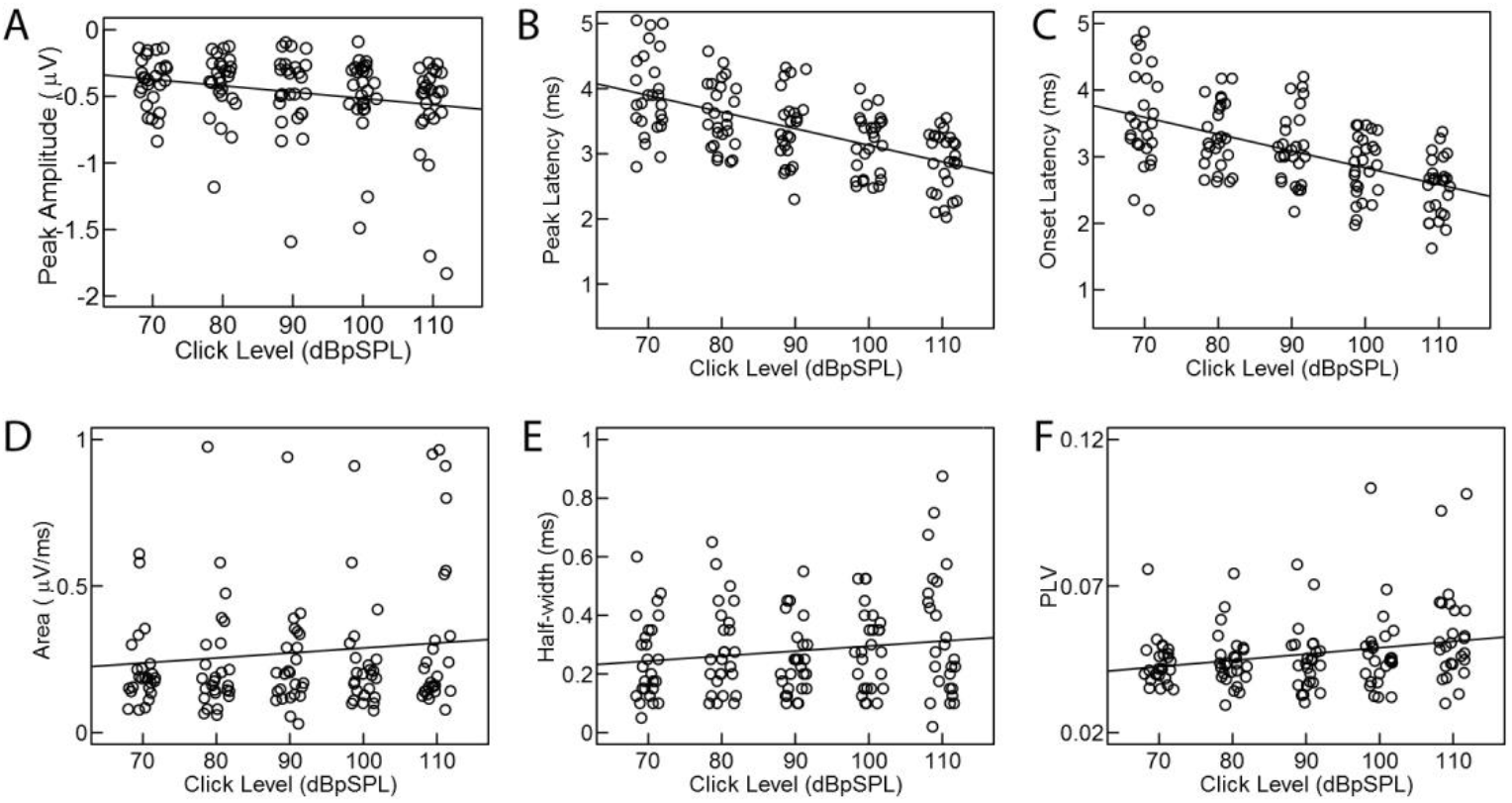
CAP metrics as a function of stimulus intensity level. With increasing level, peak-to-baseline amplitude increased (became more negative) **(A)**, peak and onset latencies decreased (**B** and **C**), and half-width increased (**E**), as expected. Although not significant, Area increased with increasing stimulus level (**D**). PLVs increased with increasing level (**F**), indicating improved synchrony of N1. Individual data points were randomly jittered around each intensity level (70, 80, 90, 100, and 110 dB pSPL) for clarity in display. Each dot represents data from a single participant and the solid line represents the regression of the variables by intensity level.

### 3.2 CAP metrics: Individual differences

Studies of the CAP have suggested that the response amplitude of N1 is indicative of the quantity and integrity of AN fibers. We hypothesized that the relationship of CAP metrics at low and high intensity levels will vary across individuals and levels based on individuals’ ability to recruit additional AN fibers with increasing intensity. Therefore, we examined how each metric related to N1 amplitude at a lower (80 dB pSPL) and a higher (110 pSPL) intensity level. Stronger phase locking (PLV) was associated with larger N1 responses (Figure 6) at lower (r= -0.68, p=0.001, 95%CI = [-0.86, -0.30]) and at higher intensity levels (r= -0.85, p<0.001, 95%CI = [-0.96, -0.27]). Given that phase locking is fundamental to the generation of an averaged event-related potential (ERP), PLV may explain some of the variance in ERP amplitudes throughout the auditory system. Consistent with this interpretation we have previously reported associations between the amplitude of the auditory cortical evoked response and PLV (Harris et al. 2014). Our current results provide a novel metric to assess PLV at the level of the AN and therefore disambiguate differences attributed to synchrony from other contributing factors, such as the recruitment of additional AN fibers.

**Figure 6.**
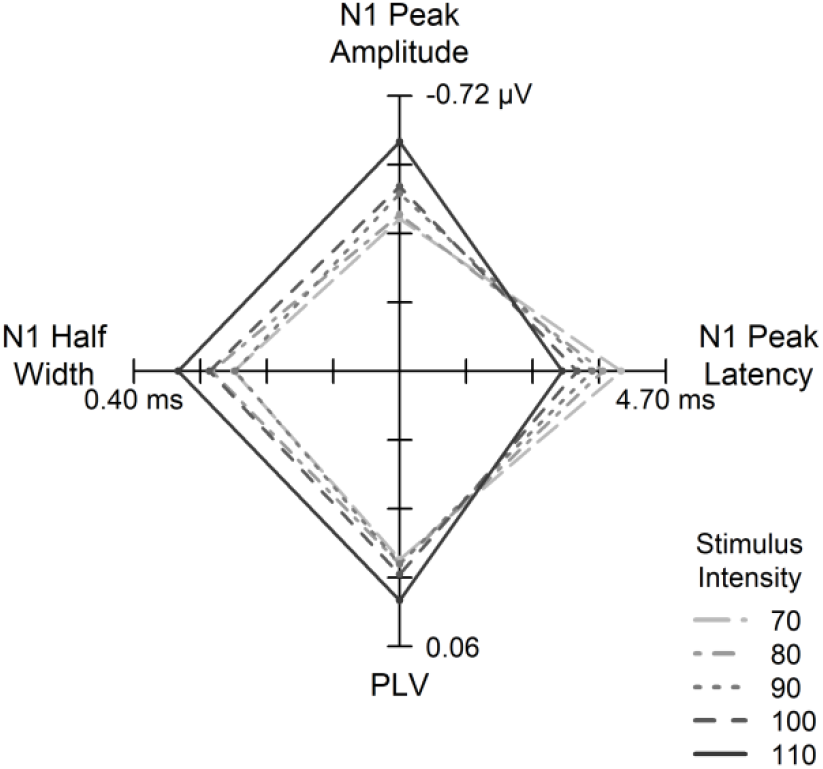
Association between CAP metrics across intensity level. A star plot with four axes illustrating the multivariate pattern of intensity-dependent changes across N1 peak amplitude, N1 peak latency, N1 half width, and PLV. The coordinates of the origin are 0, 0. Higher stimulus intensity resulted in broader N1 half widths, higher N1 peak amplitudes, earlier N1 peak latencies, and increased phase-locking. Different shading and line types indicate the stimulus intensity associated with each combination of N1 values.

Not surprisingly, CAP area was significantly negatively associated with peak amplitude at low(r=-0.76, p<0.001, 95% CI = [-0.93, -0.69]) and high (r=-0.68, p=0.001, 95% CI = [-0.94, -0.27]) intensity levels (results not shown). However, CAP area was the only variable tested that did not show a significant change with signal level (Figure 4D), likely due to increased variability in this measure. Area is measured from the beginning to the end of the N1. The offset of the N1 is dependent on the following P2 response and shape of the CAP response function. The P2 response is more variable within and across participants than the N1 (data not shown) and may limit the clinical utility of this area measurement.

As described earlier, CAP amplitude increased (became more negative) and latency decreased with increasing stimulus intensity. However, these changes were not uniform across participants or CAP metrics, with some participants and metrics showing steeper rates of change than others. We hypothesized that these individual differences in the rate of change across variables may reflect differences in the integrity of the AN, specifically the recruitment of low-SR fibers. Latency (peak and onset) and amplitude metrics were independent of one another at lower intensity levels (p>0.05). Larger peak amplitudes were associated with *prolonged* N1 *peak* latencies (r= -0.64, p=0.003, 95%CI = [-0.82, -0.33]) and wider (half-width) responses, at higher levels (r= -0.75, p<0.001, 95%CI = [-0.90, -0.27]) (Figure 6E, F). Peak amplitudes were not significantly correlated with *onset* latencies at either level (not shown, p>0.05), suggesting that larger peak amplitudes with prolonged peak latencies are not simply the result of delayed N1 responses. The amplitude and latency of the response to the onset of a stimulus is determined by the number and temporal pattern of responding neurons; their timing is calculated as the first spike latency. First spike latencies decrease with increasing SR and characteristic frequency(Heil and Irvine 1997). Therefore, prolonged peak latencies and larger amplitudes at higher but not lower levels are consistent with the recruitment of low-SR fibers.

In addition to reductions in maximum amplitude, physiological findings using the CAP recorded from the AN in animals show shallow slopes of input-output functions in those animals with a loss of AN fibers (Hellstrom and Schmiedt 1990; 1991; Schmiedt et al. 1996). Furthermore, while absolute differences in CAP amplitude may stem in part from several factors unrelated to the AN, including placement of the electrode, tissue conductivity, and electrode resistance, differences in slope normalizes these effects. We observed substantial differences in the slope of the CAP amplitude-intensity function across participants. We hypothesized that increased neural synchrony across trials (PLV) and/or the recruitment of low-SR fibers would contribute to individual differences in the increase in CAP amplitude at high intensity levels. We used multiple linear regression and model testing to determine the extent to which PLV and N1 peak latency (110 dB pSPL) uniquely contributed to the growth of N1 amplitude with increasing intensity. Both PLV (p=0.002) and peak latency (p=0.008) remained strong predictors of N1 amplitude growth (Multiple R^2^ =0.58, p<0.001). Model testing showed that including both PLV and peak latency significantly improved model fit compared to a model where only PLV was a predictor (F (1, 23) =8.42, p=0.008) or a model where only peak latency was a predictor (F (1, 23=12.07, p=0.002) (Table 1). Moving forward, assessing the contribution of synchrony and latency shift with growth of N1 amplitude with increasing intensity may help characterize the underlying pathology present in patient populations or across age groups.

**Table 1.** PLV and N1 latency associations with N1 amplitude growth

| **Model 1** | $\beta$ | $t$ score | $p$ value |
| --- | --- | --- | --- |
| PLV | -0.30 | -4.27 | $p < 0.001$ |

Multiple R squared = 0.43, $p < 0.001$
| **Model 2** | $\beta$ | $t$ score | $p$ value |
| --- | --- | --- | --- |
| N1 peak latency | -0.009 | -3.72 | $p = 0.001$ |

Multiple R squared = 0.38, $p = 0.001$
| **Model 3** | $\beta$ | $t$ score | $p$ value |
| --- | --- | --- | --- |
| PLV | -0.23 | -3.48 | $p = 0.002$ |
| N1 peak latency | -0.006 | -2.90 | $p = 0.008$ |

Multiple R squared = 0.59, $p < 0.001$

### 3.3 Relationship to pure-tone thresholds and the acoustic reflex

All participants had thresholds ≤20dB HL from 250-8000 Hz (i.e., within clinically normal limits). Although some degree of variability was evident across participants, pure-tone thresholds at all frequencies were not significantly associated with neural metrics or results of the ITD digit segregation task (p>0.05). With respect to the acoustic reflex, when sounds reach a certain high intensity level (which varies across people and stimuli), a reflex is initiated with an efferent (ipsilateral and contralateral) contraction of the stapedius muscle. This reflex restricts movement of the tympanic membrane, dampening the acoustic signal delivered to the cochlea. Therefore, individual differences in CAP metrics at low versus high intensity levels could stem from differences in the level at which the acoustic reflex is elicited. However, none of the six participants tested exhibited an acoustic reflex within the range of levels used in the current experiment (70-110dBpSPL) and measured acoustic reflex thresholds ranged from 115 to more than 120 dB pSPL. Furthermore, complete lesion of cochlear efferent nerves in animal models fails to alter peak latencies of wave I of the ABR (equivalent to the CAP)(Maison et al. 2013), arguing against a contribution of the acoustic reflex to level-dependent changes in CAP metrics.

### 3.4. Relationship between suprathreshold temporal processing and CAP metrics

Overall, ITD digit segregation improved with increasing ITD (F (5, 65) =8.68, p<0.001), but large individual differences in performance were observed. Errors generally arose because listeners incorrectly reported the digit from the masking stream that co-occurred with the target digit and not from omissions or lapses in intelligibility that would result in a misidentification. As results of previous studies have suggested, this pattern of errors is more likely to stem from individual differences in sensory encoding that make it difficult to spatially resolve the target from the masker than from lapses in attention or working memory (Bharadwaj et al. 2015). To evaluate the relative contribution of AN function versus traditional metrics of audibility (pure-tone average) we entered the data into generalized linear regression analysis. The data were modeled using a linear mixed-effects approach with a random effects term for participants. ITD, individual N1 amplitude slopes, and pure-tone average (0.5, 1, 2, and 4 kHz) were entered as explanatory variables, and correct trials/ITD level as the dependent variable. The coefficient for the pure-tone average term was not significant (p=0.86). The ITD condition (t (108) = 5.09, p< 0.001) and N1 amplitude slope (t (22) = -2.26, p= 0.03) were significant predictors of performance. Interactions between N1 amplitude slope and ITD were not significant (p>0.05) (Table 2). Post-hoc Pearson product correlations showed that participants with steeper N1 amplitude slopes (more negative) had better performance on the ITD digit segregation task (r= -0.53, p=0.02, 95%CI = [-0.79, -0.22]) (Figure 7). ITD digit segregation is a difficult task with obvious demands on selective attention. However, our results suggest that individual differences in sensory processing, occurring as early as the AN, may also explain a significant portion of the variance in performance.

**Figure 7.**
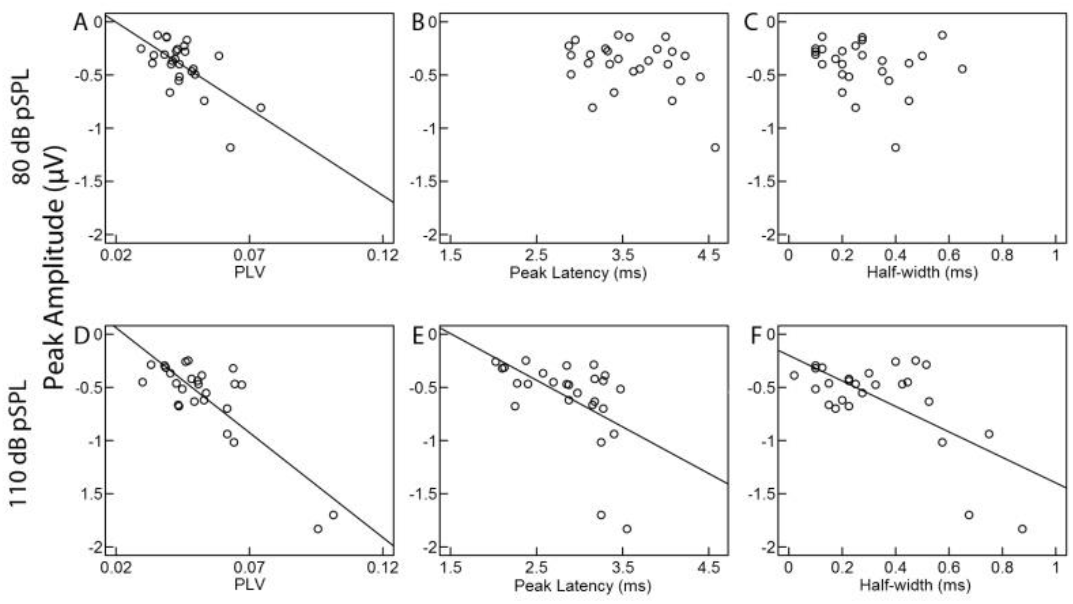
Scatter plot of N1 peak amplitude as a function of CAP metrics and PLV, at 80 dB pSPL (top) and 110 dB pSPL (bottom). Peak amplitudes increased (became more negative) as PLV increased at low (**A**) and high (**D**) levels, consistent with more precise neural synchrony across stimulus trials generating larger N1 peak responses. Unexpectedly, larger peak amplitudes were also associated with longer N1 *peak* latencies and wider (half-width) responses, but only at the higher level (**E** and **F**). Peak amplitudes were not significantly correlated with *onset* latencies at either level (data not shown, p>0.05), suggesting that larger peak amplitudes with prolonged peak latencies are not simply the result of delayed N1 responses. The solid line represents the regression line between the variables.

**Table 2.** ITD level, N1 amplitude, and pure-tone average associations with ITD digit discrimination

| **Model 1** | $\beta$ | $t$ score | $p$ value |
| --- | --- | --- | --- |
| ITD level | 0.01 | 5.09 | $p < 0.001$ |
| N1 amplitude slope | -0.02 | -2.26 | $p = 0.03$ |
| Pure-tone average | -0.003 | 0.18 | $p = 0.86$ |

Marginal R squared = 0.19
Conditional R squared = 0.91

### 3.5 Noise Exposure History

Noise history questionnaires indicated that 60% of participants had a positive history for one or more of the noise exposure categories and were included in the “exposed” group (see methods), with the most common positive response to loud music (N=10) and guns (N=8). In contrast to previous studies, the likelihood of a positive noise history did not significantly differ with sex of the participant (p > 0.05). Although pure-tone thresholds ranged from -5 to 20 dB HL, pure-tone thresholds did not significantly differ between the two groups at any frequency (p>0.05). Compared to the “non-exposed” group, the “exposed” group had significantly shallower N1 amplitude-intensity slopes (t (23) =2.23, p=0.03; Figure 8) and narrower N1 responses (t (23) =-2.09, p=0.048) with earlier peak N1 latencies (t (23) =-2.25, p=0.03). These effects were significant based on bootstrap tests with 10,000 permutations. There were no other significant differences between noise-exposure groups, including PLV (p=0.16).

**Figure 8.**
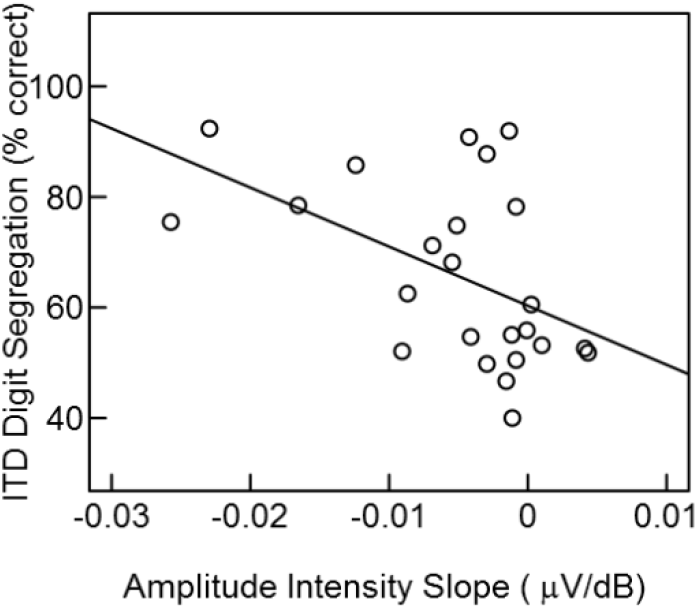
Scatter plot showing individual differences in average performance (percent correct across ITD) on the ITD digit segregation task as a function of the slope of the CAP amplitude-intensity function. 24 out of the 26 participants completed the ITD digit discrimination task. Those participants with the steepest N1 amplitude-intensity functions had the best ITD digit segregation performance.

**Figure 9.**
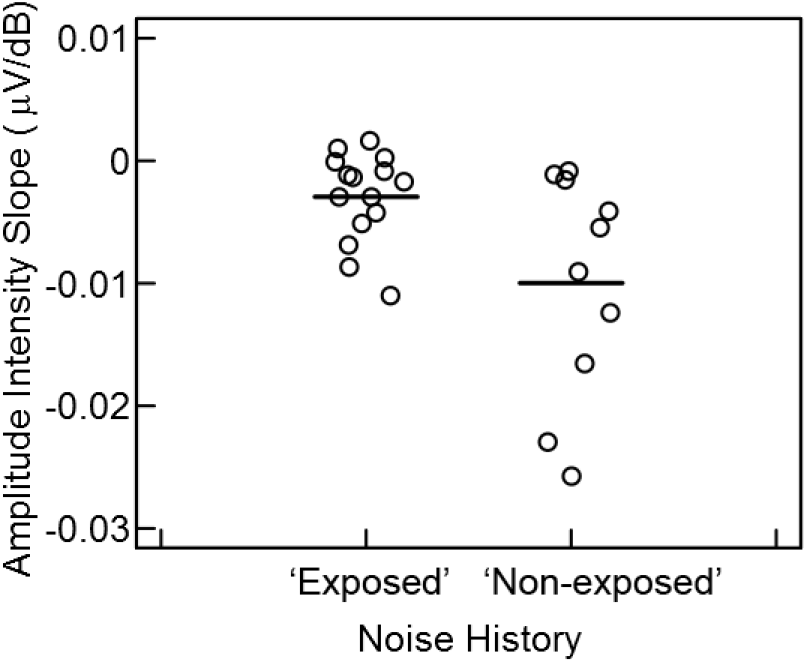
Slope of the CAP amplitude-intensity function for participants in exposed and non-exposed noise history groups. Participants in the “exposed” group exhibited significantly less N1 amplitude growth with increasing intensity (shallower functions) than the non-exposed group (p < 0.05). Horizontal lines indicate the mean slope of each group.

## Discussion

Growing evidence from animal and human studies suggests that a loss or dysfunction of AN fibers, particularly those with low-SRs, contributes to suprathreshold auditory processing deficits (Kujawa and Liberman 2009; 2015; Liberman et al. 2015; Schmiedt et al. 1996). The goal of the current experiment was to develop and validate new metrics from the CAP to better characterize AN function in humans and reveal the contributions of neural synchrony and slower AN, such as those with low-SR fibers. This was the first study to use a single-trial approach to assess the CAP response, calculating neural synchrony across trials. Our estimates of neural synchrony from the PLV of the CAP were assessed with good reliability in all participants and increased with increasing signal level and peak amplitude, suggesting greater synchrony of neural discharges at higher signal levels. These results are consistent with those from single-unit studies in laboratory animals that have shown a reduction in neural jitter with increasing signal level (Miller et al. 1999).

Although direct counts of low-SR and high-SR fibers are not currently possible in humans, we attempted to estimate the contribution of low-SR fibers by 1) using high sound levels, in which the contribution to the overall response of low-SR fibers is greater than at lower levels; and 2) examining the growth of amplitude and the width of the CAP across these higher levels. Even in our sample of younger adults with normal pure-tone thresholds we found large individual differences in CAP response metrics and suprathreshold temporal processing. We found that neural synchrony and longer peak latencies independently contributed to the growth of the CAP amplitude with increasing intensity. By examining differential measures across intensity, we demonstrated that associations between amplitude and prolonged peak latency were stronger at higher levels where the amplitude growth is dependent on the contribution of low-SR fibers, relative to lower levels. Although level-dependent increases in synchrony result in larger CAP responses, these changes do not account for level-dependent increases in N1 response latency and response width. Our examination of the onset of the response and the width of the CAP suggest that these differences in peak latency are not related to a delay or prolongation of the response but instead represent a wider response, consistent with the recruitment of slower AN fibers. Together, these complementary metrics may provide a means to assess and differentiate the underlying neural pathologies of hearing loss and aging, and better understand the interrelated roles of neural dyssynchrony and AN degeneration and loss.

Of importance to the current paper is the extent to which low-SR AN fibers contribute to the generation of the CAP response. A recent paper by Bourien et al. (2014) examined the contribution of AN fibers to the CAP; increasing doses of ouabain were delivered to the round window of Mongolian gerbils and guinea pigs. At 80 dB SPL, a selective loss of low-SR fibers appeared to have no effect on the CAP response. In contrast, experimental manipulations, both noise and furosemide, that resulted in a loss or dysfunction of low-SR fibers, resulted in robust changes in Wave I of the ABR, including increased thresholds and decreased response amplitudes(Fernandez et al. 2015; Lang et al. 2010; Mehraei et al. 2016; Sergeyenko et al. 2013). The contribution of low-SR fibers to the CAP has also been demonstrated in the recovery of the CAP response from prior stimulation, both in animals where the distribution of low-SR fibers was estimated (Schmiedt et al. 1996), and in humans (Murnane et al. 1998). Importantly, the study by Bourien et al. (2014) assessed the contribution of low-SR fibers with stimuli presented at 80 dB SPL, and at this intensity level our results are consistent with this finding, showing little to no association between response width or peak latency and amplitude. finding, showing little to no association between response width or peak latency and amplitude. Associations among metrics do not reach significance until higher intensity levels, consistent with the higher thresholds of low-SR fibers, and known saturation rates of high SR fibers (50- 80 dB SPL) (Sachs and Abbas 1974).

In animals, cochlear synaptopathy and deficits in AN function can be identified via the suprathreshold amplitudes of the CAP response or wave I of the auditory brainstem response (ABR) (Fernandez et al. 2015; Sergeyenko et al. 2013). In humans, the use of CAP amplitudes as a marker of AN integrity is complicated by factors unrelated to AN integrity, including electrode location and resistance, head size, etc. To minimize these confounds we adopted a multi-metric and differential approach, examining the interaction of variables at multiple intensity levels. This approach reduces is based on the assumption that the impact of low-SR fibers would be greater at higher than lower intensities. Recently, several additional methods have been proposed for assessing AN loss and function in humans. Liberman et al. (2016) suggested using the ratio of the summating potential (SP), generated by the inner hair cells (IHC) in the cochlea, to the CAP N1, based on the assumption that ‘hidden hearing loss’ and cochlear synaptopathy would affect the AN (i.e., CAP response) while leaving the IHCs intact (i.e., the SP). While they did indeed find a significant difference in the SP/CAP ratio between their ‘noise-exposed’ and ‘non-exposed’ groups, these differences were driven by differences in the SP and not the CAP, complicating interpretation. The click stimulus, 200 Hz filter, and presentation rate in the current study were selected to optimize the CAP response; therefore we were unable to compare our metrics to the SP/CAP ratio because the SP could not be recorded reliably in several participants. In addition to measures from the AN, several studies have advocated for the use of brainstem measures, including the frequency following response (FFR) and wave V of the ABR, to identify deficits in AN function (Bharadwaj et al. 2015; Mehraei et al. 2016). However, these responses are generated at the level of the brainstem and individual differences in both central and AN function could contribute to the response. This limitation is of particular importance when examining different experimental or patient populations, as both age and experience are known to affect central neural function (Clinard and Tremblay 2013; Clinard et al. 2010; Krishnan et al. 2016; Wong et al. 2007). Therefore, changes observed in these responses could be a consequence of a combination of AN and central factors. In contrast, our AN measures should reflect AN function across study and clinical populations, particularly after controlling for peripheral differences using pure-tone thresholds or otoacoustic emissions (Liberman et al. 2016).

### Relationship to temporal processing

Similar to previous studies employing the digit stream segregation identification task (Bharadwaj et al. 2015); we found large variations in perceptual ability, even across younger adults with normal pure-tone thresholds. This task was designed to limit non-sensory factors that affect selective attention demands during speech recognition task performance in order to focus on the degree to which sensory processing differences impact performance. The ability to differentiate the two digit streams was dependent on the use of small ITD cues, thereby testing the fidelity of the temporal resolution in the auditory system. While individual differences in suprathreshold AN activity were predictive of performance, performance was unrelated to small differences in hearing thresholds. These results support the hypothesis set forth by Bharadwaj et al. (2015) that the fidelity of temporal feature encoding, even at early levels of the auditory system, may contribute to ITD stream segregation. The synchronous neural firing patterns of AN fibers are critical for encoding time-varying acoustic cues, particularly when speech is presented in the presence a competing speech stream. A loss or dysfunction of the AN may result in an stochastic undersampling of these speech streams, (Lopez-Poveda and Barrios 2013) contributing to difficulties in segregating the digit streams.

### Effects of noise on CAP responses

Noise exposures, even moderate exposures, are neuropathic and can lead to changes in the CAP response, including decreased peak latencies and decreased CAP amplitudes (Lichtenhan and Chertoff 2008; Maison et al. 2013). Even when the noise is carefully calibrated, large individual differences are often observed. Recreational noise exposure is more difficult to quantify, but some estimates suggest that recreational exposure often exceeds 100 dB SPL (Clark 1991). Noise history questionnaires used to assess noise exposure in humans are subjective in nature, particularly when assessing exposure to ‘loud’ sounds, and are subject to recall bias. Despite these limitations, we observed significant differences between ‘exposed’ and ‘non-exposed’ groups, consistent with a loss or inactivity of AN fibers. These results are consistent with models developed by Lichtenhan and Chertoff (2008), suggesting that following a noise exposure, even one that only results in a temporary elevation in hearing thresholds, latencies of the CAP decrease and response is narrower due to fewer AN fibers contributing to the CAP response. Our results are also consistent with recent reports that have identified small but significant associations between self-reports of noise exposure and the CAP response(Bramhall et al. 2017; Liberman et al. 2016; Stamper and Johnson 2015).In summary, we provide new metrics that describe the CAP response function to characterize AN function *in vivo*. Future studies will be designed to confirm underlying mechanisms and apply these CAP metrics and behavioral measures to characterize age- and hearing-loss-related changes in AN function. Together, these complementary metrics may provide a means to assess and differentiate the underlying neural pathologies of human presbyacusis and better understand the interrelated roles of neural dyssynchrony and AN degeneration and loss. Our results are consistent with a role of low-SR fibers and changes in low-SR fiber populations that can occur with noise exposure and age. However, differences observed in AN function among younger adults may simply represent typical developmental variability/experience or genetic/epigenetic factors not evaluated in the current study, rather than effects of cochlear synaptopathy. More research is needed to rule out these contributing factors and confirm the associations between CAP metrics and AN function. One approach to the validation of these measures is the use of computational models in humans or animal models. In animal models of cochlear synaptopathy, substantial lesions in the AN resulted in near normal thresholds. However, differences in N1 amplitude of the CAP at suprathreshold levels, and model estimates of the number of AN fibers significantly correlated with measures of surviving AN area (Earl and Chertoff 2010).

## Acknowledgements

We thank Dr. Richard Schmiedt for his early review and help in developing these projects. This work was supported (in part) by grants from the NIH/NIDCD (R01 DC014467 and P50 DC00422). The project also received support from the South Carolina Clinical and Translational Research (SCTR) Institute with an academic home at the Medical University of South Carolina, NIH/NCRR Grant number UL1RR029882. This investigation was conducted in a facility constructed with support from Research Facilities Improvement Program Grant Number C06 RR 14516 from the National Center for Research Resources, NIH.

